# Rapid assessment of the suitability of multi-species citizen science datasets for occupancy trend analysis

**DOI:** 10.1101/813626

**Authors:** Michael J.O. Pocock, Mark W. Logie, Nick J.B. Isaac, Charlotte L. Outhwaite, Tom August

## Abstract

- Species records from volunteers are a vast and valuable source of information on biodiversity for a wide range of taxonomic groups. Although these citizen science data are opportunistic and unstructured, occupancy analysis can be used to quantify trends in distribution. However, occupancy analysis of unstructured data can be resource-intensive and requires substantial expertise. It is valuable to have simple ‘rules of thumb’ to efficiently assess the suitability of a dataset for occupancy analysis prior to analysis.
- Our analysis was possible due to the production of trends, from our Bayesian occupancy analysis, for 10 967 species from 34 multi-species recording schemes in Great Britain. These schemes had an average of 500 visits to sites per year, and an average of 20% of visited sites received a revisit in a year. Occupancy trend outputs varied in their precision and we used expert elicitation on a subset of outputs to determine a precision threshold above which trends were suitable for further consideration. We then used classification trees with seven metrics to define simple rules explaining when the data would result in outputs that met the precision threshold.
- We found that the suitability of a species’ data was best described by (i) the number of records of the focal species in the 10% best-recorded years, and (ii) the proportion of recording visits for that taxonomic group with non-detections of the focal species. Surprisingly few data were required to be predicted to meet the precision threshold. Specifically, for 98% confidence that our Bayesian occupancy models would produce outputs meeting the precision threshold, there needed to be ≥29 records of the focal species in the 10% best-recorded years (equivalent to an average of 12.5 records per year in our dataset), although only ≥10 records (equivalent to 4.5 records per year) were required for species recorded in less than 1 in 25 visits.
- We applied these rules to regional species data for Great Britain. Data from 32% of the species:region combinations met the precision threshold with 80% confidence, and 14% with 98% confidence. There was great variation between taxonomic groups (e.g. butterflies, moths and dragonflies were well recorded) and region (e.g. south-east England was best recorded).
- These simple criteria provide no indication of the accuracy or representativeness of the trend outputs: this is vital, but needs to be assessed individually. However our criteria do provide a rapid, quantitative assessment of the predicted suitability of existing data for occupancy analysis and could be used to inform the design and implementation of multi-species citizen science recording projects elsewhere in the world.

## 1 Introduction

Biodiversity data are vital for assessing trends in species, understanding changes in ecosystems (Butchart et al., 2010; Dirzo et al., 2014) and are required for international reporting, e.g. for the Aichi biodiversity targets (Brooks et al., 2015; Díaz et al., 2019). Currently, there are major limitations in our ability to report biodiversity trends for many taxa in many parts of the world due to a lack of data (Beck, Böller, Erhardt, & Schwanghart, 2014). Citizen science is a method that can efficiently gather lots of data and so has potential to fill this data deficit, as well as providing many other benefits in empowering and engaging people (Dickinson et al., 2012). Environmental citizen science is very diverse, varying from highly structured sampling to unstructured methods, in which people can submit data when and from where they choose (Michael J. O. Pocock, Tweddle, Savage, Robinson, & Roy, 2017). These types of dataset have been called ‘unstructured’ or ‘opportunistic’ because they do not have formal protocols for data collection. Here, we call these ‘occurrence data’ because they simply contain lists of species observations from a particular date and location. Occurrence data form most of the biodiversity records available through services such as the Global Biodiversity Information Facility (GBIF, http://www.gbif.org), regional services (e.g. the National Biodiversity Network Atlas in the UK), taxonomic repositories (e.g. eBird; Sullivan et al. (2014)) and specific citizen science projects (e.g. iNaturalist).

Occurrence data are typically comprised of presence-only records of a set of species from a location. Often projects are multi-species and taxonomically-based. This means that the records can be converted into detection histories because absence on a list can be used as a non-detection: non-detections include species that were absent, species that were present but not sighted, and those sighted but not reported (Altwegg & Nichols, 2019). Occupancy analysis allows us to obtain estimates of occupancy (i.e. the probability that a species was present, whether it was seen or not), taking imperfect detection into account (Guillera-Arroita, 2017; Isaac, van Strien, August, de Zeeuw, & Roy, 2014, p. 20; MacKenzie et al., 2002; van Strien, van Swaay, & Termaat, 2013). In particular, this depends upon at least some sites being revisited within a time period over which occupancy is to be estimated (typically a year) so that detection can be estimated. Increasingly, projects encourage recorders to submit complete lists or ‘checklists’ of species (Isaac & Pocock, 2015; B. L. Sullivan et al., 2014), to help reduce bias arising from incomplete reporting of species that were sighted (Kelling et al., 2019; Szabo, Vesk, Baxter, & Possingham, 2010). Occupancy models are now considered to be a valuable approach for estimating trends in occupancy from presence-only data (Isaac et al., 2014; van Strien et al., 2013), and are widely-used for this purpose (Burns et al., 2018; Cole et al., 2019; Powney et al., 2019; Termaat, van Grunsven, Plate, & van Strien, 2015), although they have received some criticism (Kamp, Oppel, Heldbjerg, Nyegaard, & Donald, 2016).

Recent advances in occupancy modelling have made it possible to estimate long-term trends even when recording intensity is low. In particular, Bayesian approaches make it more tractable to construct complex (i.e. realistic) detection and occupancy sub-models and can share information on occupancy across years to make the trends less dependant on the vagaries of data in a single year (Charlotte L. Outhwaite et al., 2018). However, with these advances come costs. In particular, there is a high investment of time and expertise to run these Bayesian occupancy models, and with large datasets models can take several weeks to run even when using high-performance computing. It is valuable to know whether the data are suitable for analysis before investing in the expense of running these models. Being able to predict the precision of outputs before running the models could avoid wasted time and effort.

In this study, we aimed to develop simple criteria (‘rules of thumb’) to predict when occurrence data would provide trend outputs that were sufficiently precise for further consideration (e.g. in meta-analysis or biodiversity reporting). The simple criteria were therefore a form of *post hoc* power analysis for these trend outputs, but they could also be used for *a priori* screening of different data for trend analysis. In this study, we applied these simple criteria to the data for species within regions in GB, to provide a rapid assessment of the suitability of these data for regional biodiversity trend analysis.

## 2 Methods

### 2.1 Overview

We sought to undertake a rapid assessment of the suitability of multi-species citizen science occurence datasets for occupancy analysis. Our analysis was possible because of the production of a huge, recently-created dataset of occupancy model outputs, spanning 10,976 species from 34 taxonomic groups in Great Britain (Outhwaite et al., in press), from relatively well-recorded taxonomic groups (e.g. bees and dragonflies) to those which are sparsely recorded (e.g. centipedes and fungus gnats). 5,293 of these trend outputs (comprising all species with >50 records) have already been published (Outhwaite et al., 2019).

In this study, our approach was three-fold. Firstly we know that trend outputs varied in their precision (Fig. 1), but there was no accepted measure of when the precision of outputs was unacceptably low. We used expert elicitation on a subset of outputs to determine a precision threshold above which trends were suitable for further consideration. Secondly we developed seven metrics to describe the occurrence datasets. We used classification tree analysis to define simple criteria predicting when the data would meet the precision threshold. Thirdly we applied the simple criteria to species records in GB regions. This was to assess taxonomic variation in the regional coverage of recording in GB.

**Fig 1.**
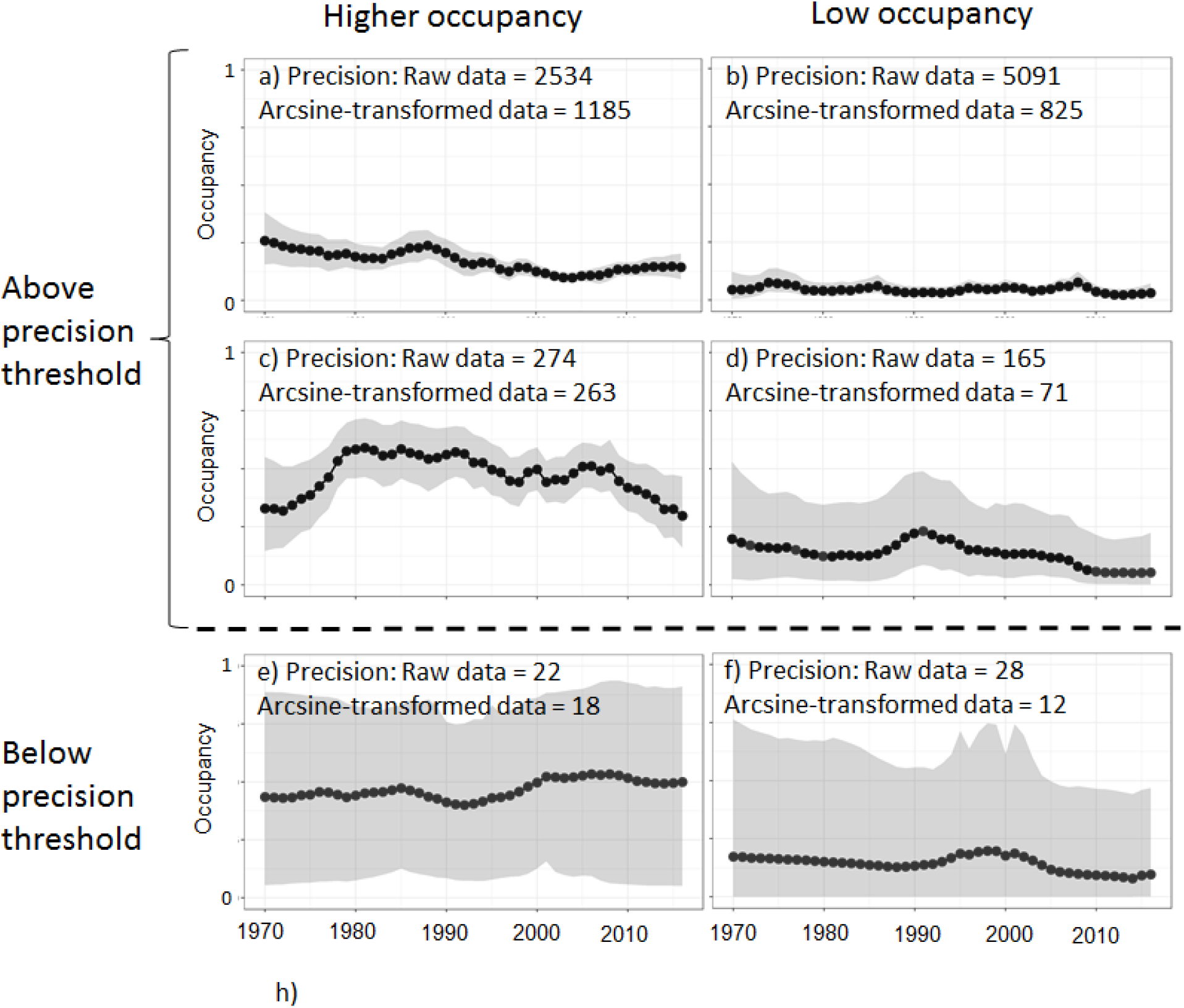
Examples of occupancy trend outputs showing species with higher occupancy (mostly >0.25; a, c, e) and low occupancy (<0.25; b, d, f). The trends are grouped by whether they met the precision threshold (see Results; specifically, this was defined according to the precision of the arcsine square root-transformed posteriors of the Bayesian occupancy analysis). The species shown are all bees: (a) *Bombus bohemicus*, (b) *Ceratina cyanea*, (c) *Nomada marshamella*, (d) *Sphecodes ferruginatus*, (e) *Lasioglossum sexnotatum*, (f) *Andrena vaga*, with the data available in Outhwaite et al. (in press).

Our results are therefore based on the specific pattern of biodiversity recording by volunteers in GB and the formulation of the Bayesian occupancy models employed by Outhwaite et al (2019).

However, we expect that the general lessons will be broadly applicable to similar types of analysis (i.e. occupancy analysis) on similar datasets (i.e. from multi-species citizen science projects) elsewhere in the world.

### 2.2 Description of datasets

Here we used trend outputs from Bayesian occupancy analyses that had already been run for a total of 10,967 species from 34 multi-species recording schemes (hereafter called ‘projects’; median species per project = 169; range = 9-1002); many of these are publically available (C. L. Outhwaite et al., 2019), although outputs from species with less than 50 records have not been released (Outhwaite et al., in press). The Bayesian occupancy analyses were run over the 47 year time period under consideration (1970 to 2016). Recording intensity has increased over these four decades, so in order to define simple rules that are appropriate in different situations, we also used the last ten years’ data in our analysis. Specifically, rather than re-running the models, we used the trend output and data metrics from the last ten years separately from the 47 year trends. Although the data and outputs for the 10 year period are a subset of the 47 year period, they could vary considerably in recording intensity or species trend and so we treated them as if they were independent. Each species was therefore included twice: for the 47 year and ten-year trends.

The data supporting these analyses came from national biological recording schemes in Great Britain, supported through the UK’s Biological Records Centre (Pocock, Roy, Preston, & Roy, 2015). There are 85 schemes in Great Britain: they are defined by their taxonomic scope (e.g. Odonata, butterflies, bryophytes etc.), mostly led by volunteers, and since most records are submitted by people voluntarily, we regard the schemes as a form of ‘citizen science’, although recorders do need sufficient identification skills to participate. (Environment Agency staff provided data for some freshwater groups; but we do not distinguish their records because they were included in the recording scheme’s dataset; Appendix S1). Records are verified by experts (mostly volunteers) before being accepted as correct and included in datasets. Here, we regarded each recording scheme (or taxa within recording schemes, where these are recorded differently, e.g. bees, wasps and ants) as an individual ‘project’, and used data from 34 projects (Appendix S1). We treated data from each project as independent from all others.

The datasets comprise lists of species recorded in ‘visits’. A ‘visit’ is a list of one or more species reported at a specific place (here, a 1km GB Ordnance Survey grid square) and time (here, a specific day). Therefore visits to the same 1km grid square on the same date are considered to be a single visit, even if they happened to be from different recorders or different locations within the grid square. Records that were not specific to species, a day or to a 1km grid square were excluded. When a species within the taxonomic remit of each project was not reported on a visit, this was treated as a ‘non-detection’ (Altwegg & Nichols, 2019; Guillera-Arroita, 2017). For a complete description of the data and the occupancy model used, see Outhwaite et al. (in review).

### 2.3 Defining the precision threshold

We used expert elicitation on a subset of occupancy trend outputs to determine a precision threshold. This precision threshold acts as a filter: outputs failing to meet the threshold are deemed too imprecise to be useful, whereas outputs that meet this threshold are worth further consideration for analysis or reporting. The sites were visited according to the recorders’ choice and not according to a statistical design. This means that meet the precision threshold does not mean that outputs are accurate or representative (Mackenzie & Royle, 2005): outputs could be estimated precisely but wrongly.

Three experts (G. Powney and two authors: NJBI and CO) assessed 99 selected trend outputs from across the range of precision and occurrence values of our Bayesian occupancy analysis outputs. These three people undertook the assessments individually but are collaborators and so were not strictly independent. Each person was provided with images of trend outputs showing yearly occupancy estimates with 95% credible intervals and information on convergence of the parameter per year (where convergence is assessed when the Gelman-Rubin ‘Rhat’ metric (Gelman & Rubin, 1992) was less than 1.1; (Kery & Schaub, 2011)) and scored whether outputs were acceptable or not for further consideration. Subsequent discussion revealed that they classed model outputs as unacceptable when there were a large number of years with high uncertainty or failed convergence. We accepted the majority decision for each trend (the decision was unanimous for 80% of the outputs). Three variables were calculated from each trend output: (i) mean yearly precision, specifically 1/[variance of the arcsine square root transformed value of 1000 posteriors of occupancy from the Bayesian analysis], (ii) precision of the trend, where the trend was calculated from the first to last year of the analysis, and (iii) the proportion of years where Rhat was > 1.1. We used variance of arcsine square root transformed data to reduce the bias due to the boundary effect of the proportion data (Fig. S2.1; Gotelli & Ellison (2012)); this transformation has been criticized (Warton & Hui, 2011) but values near the boundary have less extreme effects (range of transformed data: 0, π/2) than logit transformation (range of transformed data: −∞, +∞). We used a decision tree in ‘evtree’ (Grubinger, Zeileis, & Pfeiffer, 2014) in R 3.5.2, with the three variables to statistically define the when trends were deemed to be acceptable for further consideration.

### 2.4 Constructing simple rules for when datasets met the precision threshold

We developed seven metrics, each describing different attributes of the data that we predicted would influence the success of occupancy analysis. These metrics were designed to be easily calculated and applicable to all species groups, regardless of species richness; they included different aspects of the dataset including records, visits and non-detections (Table 1). Prior to calculating the metrics, we undertook ‘site filtering’ by removing sites within each project where records were present in only one year (even if there had been multiple records of one or more species within that year). This site filtering had also been done before the trend analysis had been undertaken (Outhwaite et al., in press), because without this the statistical power of the occupancy analysis is reduced (see Isaac et al. (2014)). Site filtering removed an average of 32% of records from projects (range: 7-65%), with more records removed for taxa that are relatively poorly recorded (e.g. centipedes) or require specialist identification skills (e.g. bryophytes) (Appendix S1).

**Table 1.**
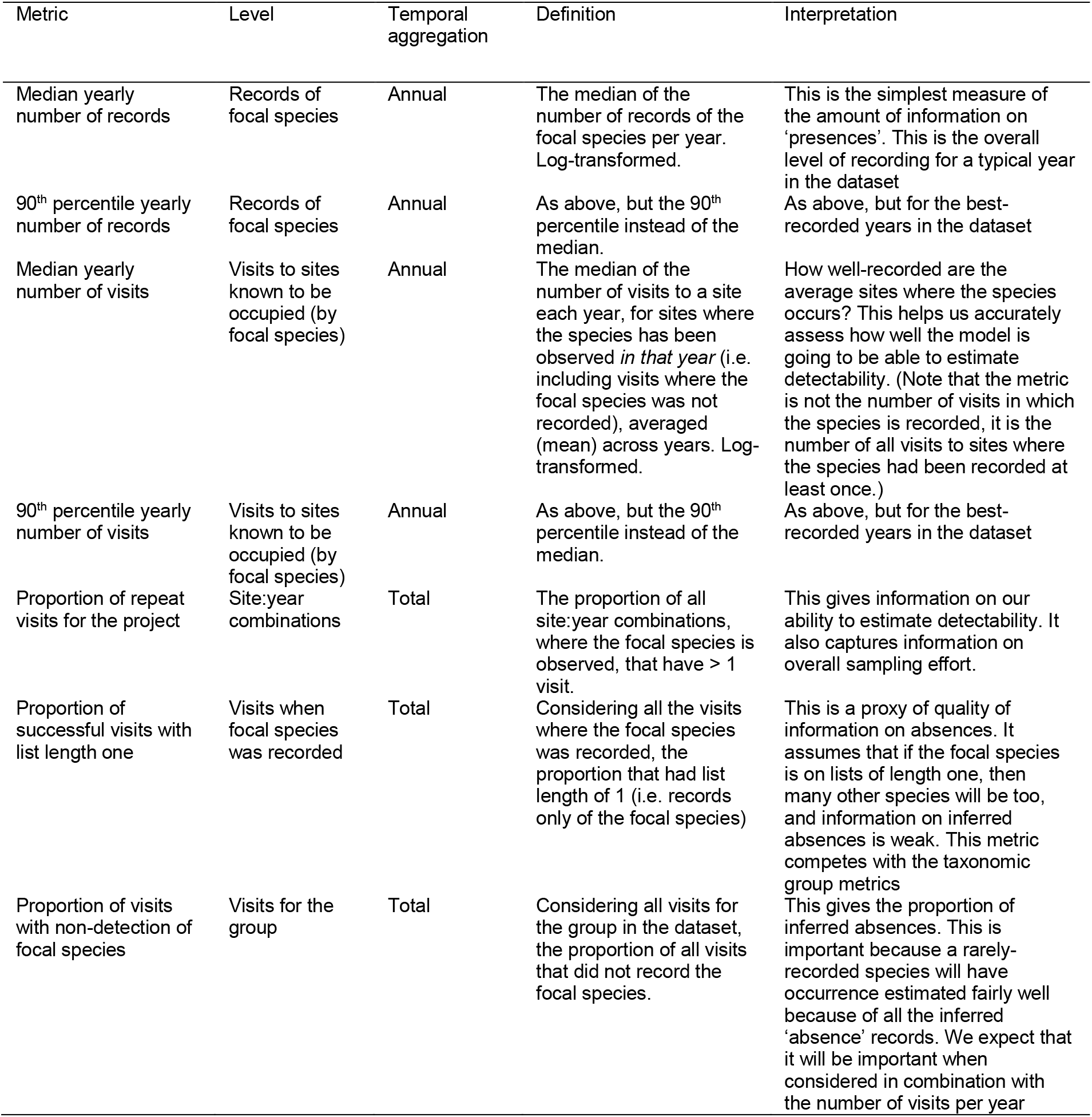
Explanation of metrics that describe variation in occurrence datasets across species and across projects.

We used decision tree analysis to construct simple criteria, based on the metrics (Table 1), to predict whether trend outputs met the precision threshold. Decision tree analysis is more appropriate than other forms of multivariate analysis because the final model retains a subset of the predictor variables and is therefore simple to interpret and efficient to apply to new datasets. Other methods, like discriminant function analysis require all metrics to make future classifications. Also, decision trees do not require predictor variables to be uncorrelated (Appendix S2). We used evolutionary classification trees, using ‘evtree 1.0-7’ in R 3.5.2 (Grubinger et al., 2014), to produce a globally optimised tree from all the data. The alternative approach, hierarchical decision trees, can mean that an optimal first-level classification can produce an overall sub-optimal tree. Decision trees were evaluated based on the balance of minimised mis-classification rate and least complexity (Grubinger et al., 2014). The depth of trees was limited to two, because initial investigation showed that decision trees with greater depth were less simple to interpret, with very little additional accuracy. We ran decision tree analysis on 31 of the datasets (7672 species; Appendix S1); we excluded moths, butterflies and lichens because of the computational demands of working with these large datasets.

We assessed the classification success of the decision tree as specificity (the proportion of trends below the threshold that were classified correctly; also called the true negative rate) and sensitivity (the proportion of trends above the threshold that were classified correctly; also called the true positive rate). Higher specificity is more conservative given the high resource costs of running models: a species meeting the criteria has a high chance that its occupancy model outputs will be above the precision threshold. On the other hand, trees with higher sensitivity mean that fewer species will discarded unnecessarily. To adjust the specificity of the decision trees, we gave data points above and below the precision threshold different weights. We therefore ran two decision trees: one tree (‘equally-weighted’ decision tree) weighed the two classes equally and so sensitivity and specificity were balanced. The second (‘high specificity’ decision tree) weighted data above the precision threshold ten times those below. These decision trees provided simple ‘rules of thumb’ criteria to assess the suitability of data for our occupancy trend analysis.

### 2.5 A rapid assessment of data suitability for regional biodiversity trends in Great Britain

We applied the ‘rules of thumb’ criteria from the two decision trees to regional biodiversity data, using the 47-year datasets for 10 967 species in 33 taxonomic groups (see Appendix S1 for details of the species included) and separating the data into 11 regions in Great Britain (using the NUTS1 statistical regions, i.e. Scotland, Wales and nine regions in England (European Union, 2016)). A small number of species had ‘no data’ for a region were because they were only recorded from sites that were visited in only one year (based on the records from the whole taxonomic group), and we had excluded these sites from analysis (see section 2.4). We calculated the proportion of species in each taxonomic group in each region that met the precision threshold according to the ‘equally weighted’ and ‘high specificity’ decision trees.

## 3 Results

### 3.1 Defining the precision threshold

The precision threshold was defined by a decision tree of the results of the expert elicitation of 99 occupancy trend outputs. The most parsimonious decision tree included only one predictor: mean yearly precision of arcsine-transformed posteriors (hereafter ‘arcsine-transformed yearly precision’), and had classification accuracy = 92%. Therefore the precision threshold, that was applied to all the trend outputs in our analysis, was defined as arcsine-transformed yearly precision = 70.4 (Fig. 2).

**Fig. 2.**
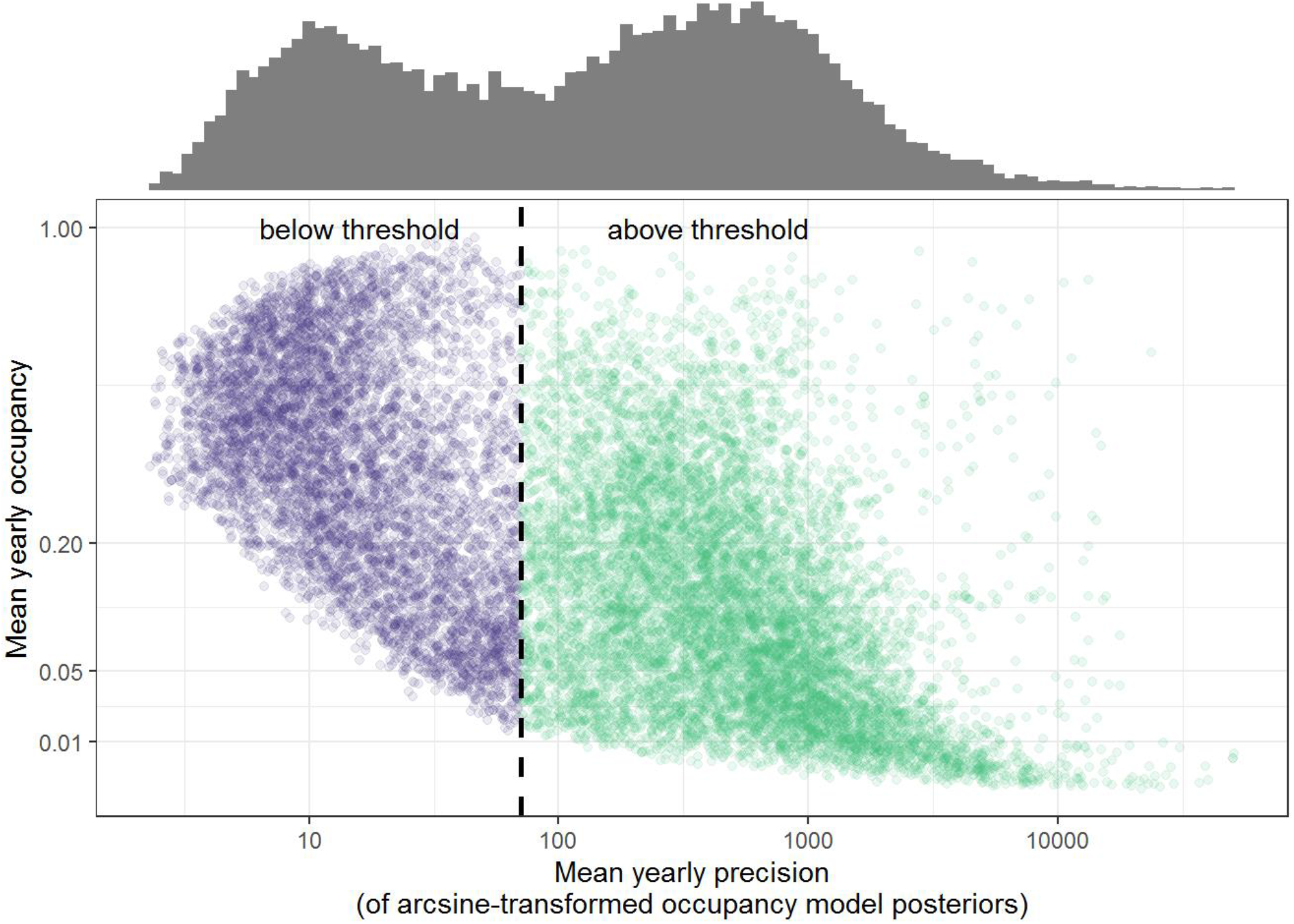
The association between mean yearly occupancy and mean yearly precision of occupancy for 15 344 trend outputs (7673 species for two time periods from 31 taxonomic groups), with the precision threshold (of arcsine-transformed model posteriors) = 70.4. For clarity the x-axis is shown on a log10 scale and the y-axis is shown on a square-root scale.

The precision threshold coincided with a trough in the bimodal frequency distribution of yearly precision for 15344 trend estimates (i.e. the 47-year and ten-year trends for the 7672 species; Fig. 2 and Appendix S3). Although we did not *a priori* expect a bimodal distribution, this provided support that occurrence datasets can be separated into those that can be modelled to give informative outputs (i.e. above the precision threshold) and those data that cannot. In our data, 57.5% of the trend outputs met the precision threshold.

For trend outputs above the precision threshold, there was a negative association between precision and occupancy (Pearson correlation *r* = −0.237; Fig. 2), indicating that species with lower estimated occupancy tend to be estimated with greater precision. This was even though we arcsine-transformed the sample of posteriors of occupancy to reduce this boundary effect. It suggests that once a species is known to have low occupancy, it is easy to be very confident that its occupancy truly is low.

### 3.2 Simple rules for predicting when a dataset will exceed the precision threshold

The decision tree analysis allowed us to construct simple criteria explaining when the data from multi-species citizen science projects are predicted to produce outputs meeting the precision threshold (Fig. 3). The equally-weighted decision tree used two data metrics for its classification: the number of focal species records for the 10% best-recorded years (90th percentile yearly number of records) and the proportion of recording visits (for that taxonomically-based project) when the focal species was not reported. This decision tree had specificity = 80.7%, sensitivity = 85.5% and overall accuracy = 83.5%. This means that 19.3% of trend outputs that met the precision threshold were misclassified, as were 16.5% of trend outputs that did not meet the precision threshold. In other words, if a dataset passes these rules of thumb we are confident (81% chance) that our occupancy modelling will lead to a trend output that will meet the precision threshold, and if it does not we are confident (86% chance) that it will not.

**Fig 3.**
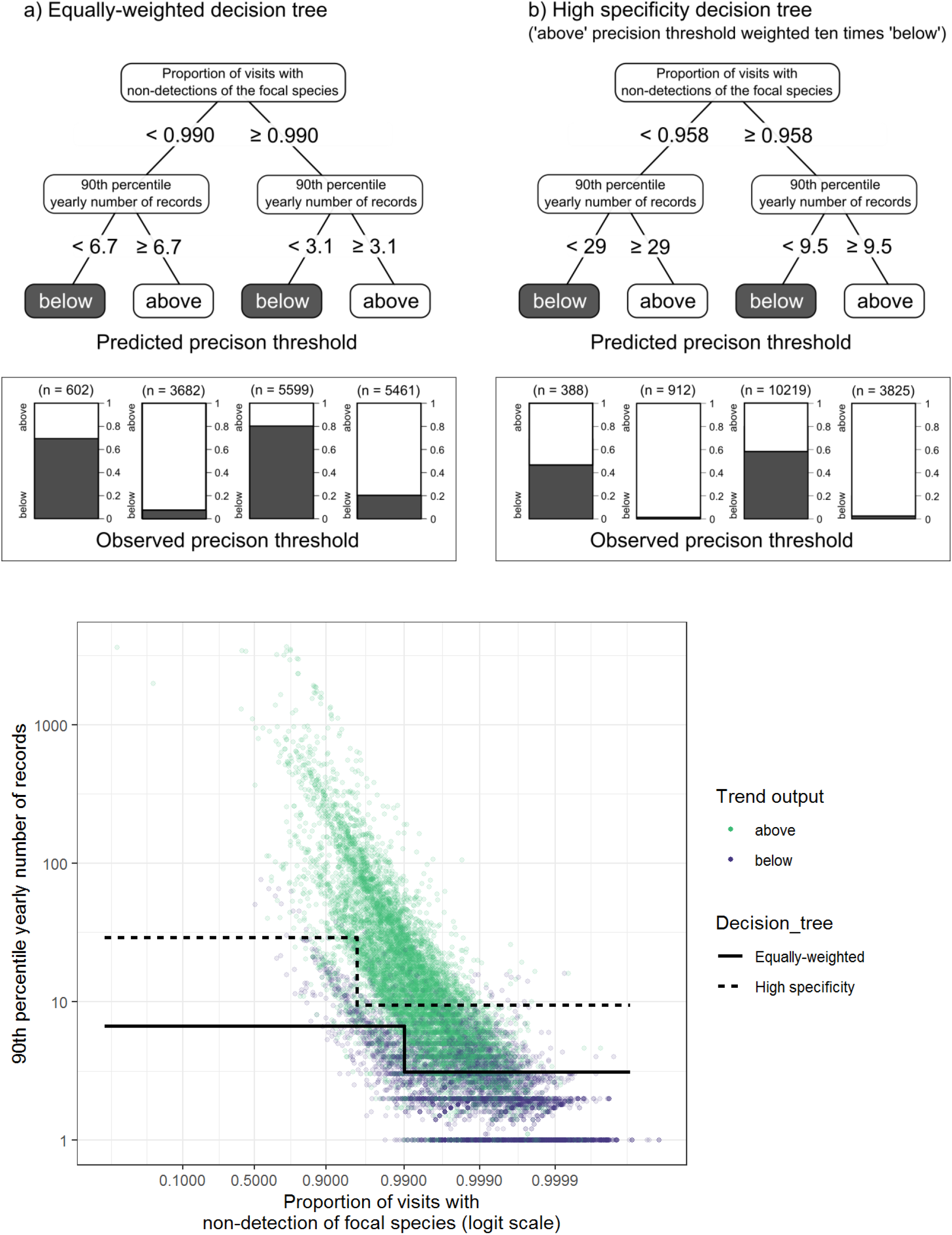
The classification trees using data metrics from 15 344 trend outputs (47-year trends and 10 year trends, treated separately for 7672 species from 31 taxonomic groups) that were (a) equally-weighted, i.e. specificity and sensitivity were balanced, and (b) high specificity, i.e. outputs above the precision threshold were weighted ten times those below. (c) This is presented graphically according to the two data metrics.

The high specificity decision tree (where the data above were weighted ten times those below the precision threshold) was similar to the equally-weighted tree, but at one node it used the metric of the median number of records rather than those for the 10% best recorded years. When the decision tree was re-run with this metric omitted there was negligible reduction in sensitivity or specificity (Appendix S3), and the result was identical in structure to the equally-weighted tree. To simplify the application and interpretation of these trees we therefore present results from this second analysis (Fig. 3). This decision tree had an specificity = 98.3%, sensitivity = 50.8% and overall accuracy = 71.0%. This means that only 1.7% of trends that met the precision threshold were misclassified, while 49.2% of trends that did meet the precision threshold were mis-classified. In other words, if a dataset passes these rules of thumb we are very confident (i.e. 98% chance) that our occupancy modelling will lead to a trend output that will meet the precision threshold, but if it does not, then it may (49% chance) meet the precision threshold anyway.

The decision trees included data metrics about both presence records and non-detections, which confirms that information from both are important for occupancy analyses (Fig. 3). The non-detection data were represented by the proportion of visits with non-detections of the focal species. In our datasets the number of records of a species in the 10% best-recorded years was significantly related to the median yearly number of records: 90th percentile yearly number of records = 1.829 × (median yearly number of records)1.058; P < 0.001 for the linear log-log relationship. This means that the 29 (or 9) records in the best-recorded years is equivalent to a median of about 14 (or 5) records per year.

In the decision trees, the number of records required in the 10% best-recorded years was lower for more rarely recorded species (Fig. 3). The definition of the more rarely recorded species varied between the two decision trees: for the equally weighted decision tree, the definition of rarely-recorded was less than 1 in 100 visits, but for the high specificity decision tree, it was less than 1 in 24 visits (Fig. 3). These more rarely recorded species actually account for the majority of species: 72.1% and 91.8%, when using the definitions from the equally-weighted and high specificity decision trees, respectively (Fig. 3). This means that most species are not recorded on the vast majority of visits, but the volume of non-detection data means that it is relatively easy to be confident that a rarely-recorded species truly occurs at low occupancy.

Although the decision trees provide clear criteria to be used as ‘rules of thumb’, it is also helpful to view these as minimum requirements. In general, more records are better because they are likely to increase precision. We found that there was a strong correlation between the minimum precision and the number of records. Based on a quantile regression to estimate the relationship of the lower limit of precision against number of records, we found that the 5th percentile of arcsine-transformed precision = 12.155 × (90th percentile yearly number of records)0.649; Fig. 4. This means that as the number of records of the focal species increased the guaranteed minimum precision of the trend outputs also increased.

**Fig. 4.**
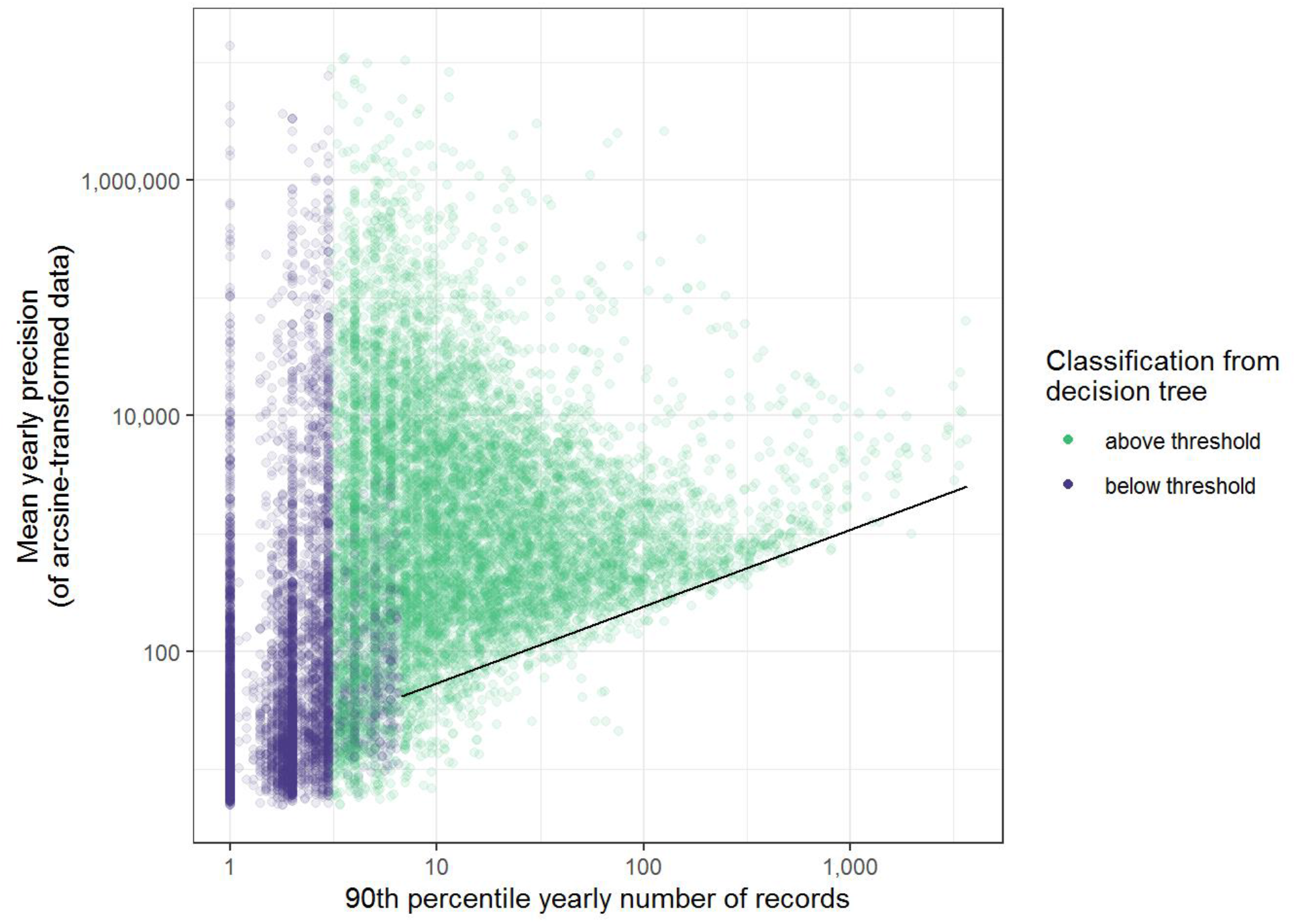
The minimum mean yearly precision is affected by the amount of data with the solid line showing a quantile regression (with the 5th percentile) for the data when 90th percentile yearly number of records > 6.7.

Our datasets were specific to the situation for biological recording in Great Britain. However, we believe that our datasets were not exceptionally unusual. The median number of visits per project per year was about 500 (first and third quartiles: 271, 1077; Appendix S1) and an average of 20% of visited sites were revisited within a year (first and third quartiles: 12%, 27%; Appendix S1). Therefore we expect that our results will be applicable to similar multi-species citizen science projects elsewhere in the world.

### 3.3 Rapid assessment of data availability for occurrence trend outputs for multiple taxa in regions of Great Britain

We considered 10 640 species from 33 taxonomic groups in Great Britain. When proportioning these by 11 regions, there were a total 62 967 species:region combinations with data. (6389 species: region combinations had ‘no data’ because the species were recorded in sites only visited once for that taxonomic group.) About one-third of the species:region combinations (20 248; 32%) were predicted to meet the precision threshold with the equally-weighted decision tree, and 8838 of these (14% of the total) were predicted to meet it with the high specificity criteria (Fig. 5). There was great variation across taxa and across GB regions in the proportion of species predicted to meet the precision threshold with our Bayesian occupancy analysis (Fig. 5; Appendix S4). However, given the huge taxonomic breadth of these projects, there are still a remarkable number of species predicted to produce trends above the precision threshold, especially when the data are aggregated at the GB-level (61.5% of the 10 640 species with data met the precision threshold with the equally-weighted decision tree).

**Fig 5.**
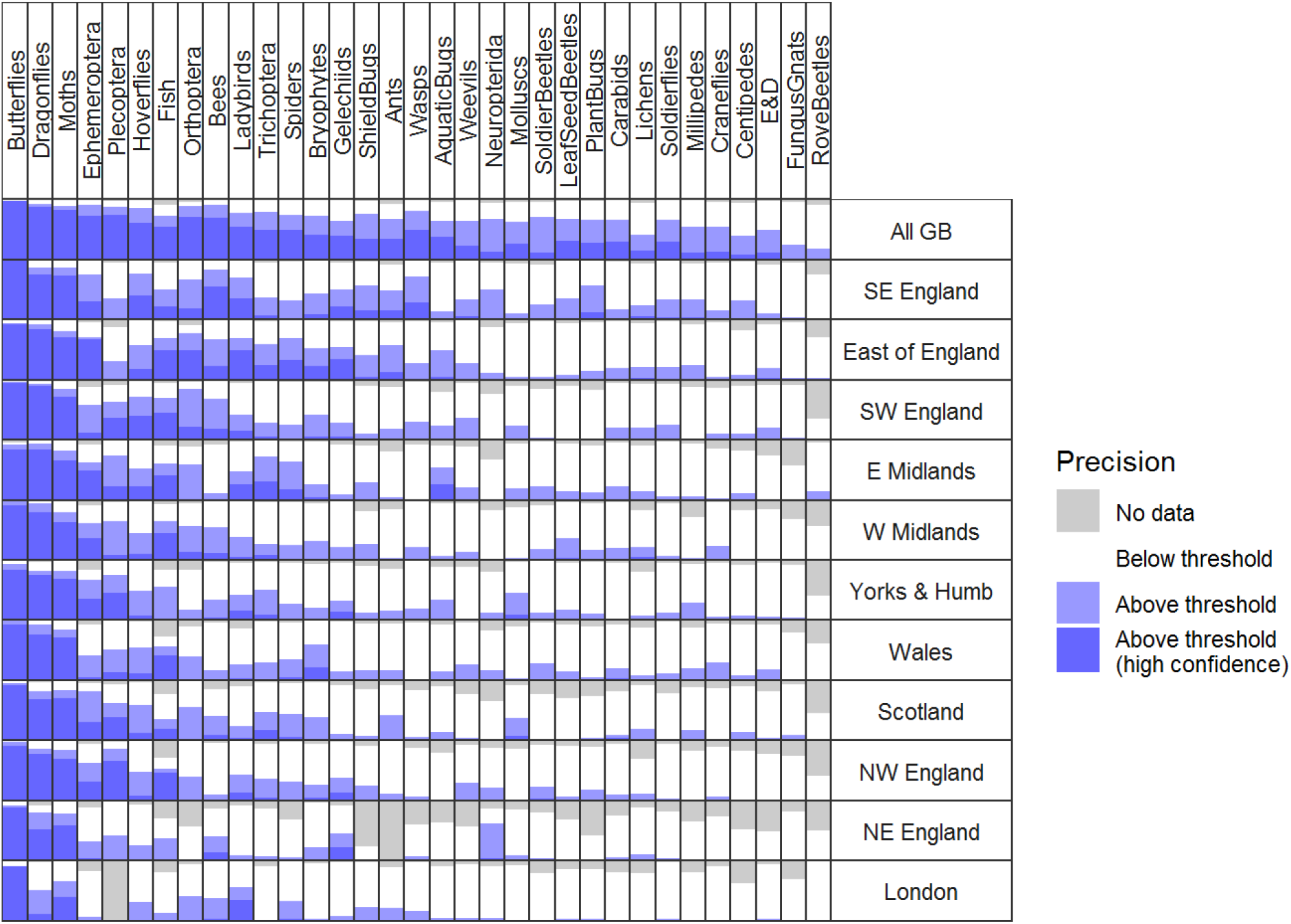
The distribution of predicted precision of occupancy trend outputs of species, with data subsets according to region in Great Britain. Data are proportion of species, using the 47-year datasets. The rows and columns are ordered by decreasing coverage.

## 4 Discussion

Here we considered a dataset of over 10 000 occupancy analysis trend outputs. We defined, post hoc and via expert elicitation, a threshold of precision: below the threshold, occupancy model outputs are unlikely to be suitable for biodiversity monitoring, whereas above the threshold, outputs can be considered for use in monitoring and research, depending on further assessment of their accuracy and representativeness. We developed indicators (‘rules of thumb’), which were based on two easily calculable metrics, to predict when occurrence data from multi-species biological recording datasets would produce outputs that meet the precision threshold. Using these indicators we were able to undertake a rapid assessment of the usefulness of occupancy models for over 60 000 combinations of species and British regions. It would have been prohibitively resource-expensive to undertake these analyses individually, so the rapid assessment is one line of evidence that allows researchers to consider the region and taxa in which it may be worth investing in future effort. Our rule of thumb criteria allow us to go beyond simply identifying variation in recorder effort (Beck et al., 2014; Meyer, Kreft, Guralnick, & Jetz, 2015; Meyer, Weigelt, & Kreft, 2016; Pocock et al., 2018) and instead allows researchers to consider when occurrence data might be suitable for occupancy trend analysis.

Due to the occupancy analysis outputs available, our study was specific to biological recording in Britain, but we are confident that the key results are applicable to the burgeoning citizen science activity elsewhere in the world. Although biological recording by volunteers has a long history in Britain (Pocock et al., 2015), there is growing interest in citizen science for biodiversity monitoring across the world (Chandler et al., 2017; Pocock et al., 2018, 2017; Sullivan et al., 2014; Theobald et al., 2015) and many projects use, or could use, taxonomically-based multi-species recording which would create the same presence/non-detection data that we considered here. We suggest that our rule of thumb indicators could be used as guidelines to identify the minimum requirements of future multi-species recording projects which have similar characteristics to our projects (i.e. a couple of hundred visits per year to sites that have been visited in more than one year, with 12-25% of those sites having more than once per year). Of course, occurrence data is not the only data that can be collected by volunteers and those developing projects in the future should carefully consider whether other approaches are better for answering the questions of interest, e.g. standardised recording of relative abundance (Pescott et al., 2019) or quantifying ecosystem function (Keuskamp, Dingemans, Lehtinen, Sarneel, & Hefting, 2013).

The rules of thumb were based on the outputs from Bayesian occupancy models that were specifically designed with weakly informative priors to be effective at estimating occupancy with sparse data, such as much biological recording data (Outhwaite et al., 2018). In these models information from detection probability are shared across sites (i.e. parameter estimates from well-sampled sites are shared with poorly sampled sites), and information on occupancy is shared across years. This makes best use of the available data, and the models perform well with simulated datasets, especially to assess long-term trends (Outhwaite et al., 2018), but it would be valuable to undertake further work on the sensitivity of the outputs to the model assumptions. Here, we took a heuristic approach in addressing this question. An alternative would have been to conduct simulations and test model outputs against truth. Simulations have been used to good effect to explore different aspects of occupancy modelling and allow a controlled approach to exploring the importance of different factors on the outputs (Banner, Irvine, Rodhouse, Donner, & Litt, 2019; Guillera-Arroita, Lahoz-Monfort, MacKenzie, Wintle, & McCarthy, 2014; Isaac et al., 2014; Welsh, Lindenmayer, & Donnelly, 2013) and provide guidelines for the design of occupancy studies (Mackenzie & Royle, 2005), although they have not been used for our specific model that shares information across sites and years. We recommend that simulations or analytical solutions can complement our heuristic approach, and it would be especially valuable to further explore the impact of the biases in sparse data on the accurate and precise estimation of occupancy trends.

When applying our rules of thumb indicators, we suggest that the high specificity decision tree is most useful for multi-species citizen science data. This is because it provides the most conservative assessment, i.e. if a dataset meets these high specificity criteria, then the output is almost certain to meet the precision threshold. This means that data need to have at least 30 records of each species in the 10% best recorded years (i.e. a median of 13 records per year in our datasets) or, for rarely-recorded species (those recorded in less than 1 in 25 visits), 10 records in the best recorded years (i.e. median of 5). Use of the high specificity criteria is at the cost of discarding potentially suitable data. We suggest that the outputs of such sparse data may be less robust anyway, and in the future this should be tested with simulations. However, if the cost of analysis is low, because expertise and computing resources are available and the data already exist, then the lower criteria from the equally-weighted decision tree would be a suitable guide.

It is important to emphasise that simply because analysis of occurrence data result in occupancy trend outputs that meet the precision threshold, this is not proof that the outputs are true: if the model is mis-specified or the data are biased then the outputs can be precise but wrong (see discussion by Dennis et al. (2019) and Kamp et al. (2016)). When considering the accuracy of the trend outputs, it is important to ask whether the trends can be considered representative. Firstly, further work is required to test the sensitivity of the trend outputs to data with uneven coverage (e.g. the well-sampled sites, which are important for estimating detectability, could be a biased subset of all sites). Secondly, expert assessment will be important, in conjunction with findings from sensitivity analysis, for case-by-case assessment of the accuracy and representativeness of the trends. In particular, the following questions are likely to be important: how are the occurrence and non-detection data distributed across the range of the species in the region of interest? and: are the visited sites representative of the region of interest? This second question is affected by biases in recorder behaviour (Tulloch, Mustin, Possingham, Szabo, & Wilson, 2013), for example leading to disproportionately more visits to nature reserves than to farmland. This assessment could be formalised as a mixed methods approach where different sources of evidence, e.g. model outputs, analysis of drivers of change, expert and public opinion, are triangulated to identify the level of agreement or areas of disagreement (Tengö, Brondizio, Elmqvist, Malmer, & Spierenburg, 2014). Thirdly, outputs can be aggregated into multi-species indicators to support policy-relevant research (Burns et al., 2018). This will not alleviate systematic biases, but it will reduce the impact of stochastic variation in results due to sparse data.

Overall, the criterion for the number of records required was surprisingly low, but these values should be considered to be minimum thresholds to be exceeded. We heuristically showed that more data from these multi-species citizen science projects is likely to improve the precision of outputs. The information provided by the presence records is combined with the information of the non-detections, but because the outputs are so reliant on non-detections it is valuable to increase the rigour of recording non-detections. This could be achieved through semi-systematic recording approaches, including complete list recording and reporting recording effort (Kelling et al., 2019). Increasing the number of visited sites is also valuable and sites must be visited in multiple years in order to be included in our Bayesian occupancy model (Outhwaite et al., 2018): simulations show that including sites visited in only one year can lead to biased estimates of occupancy (Isaac et al., 2014). About one-third of records were excluded for this reason. This means that revisiting sites that only have visits in one previous year allows both the current and the historic record to be included in the dataset, making this an activity worthy of promotion within multispecies citizen science projects.

Applying the rule of thumb indicators to the regional datasets showed the potential for cross-taxon reporting and it was certainly easier than speculatively running over 60 000 occupancy models. Assessing and explaining regional variation in biodiversity trends is important for conservation (Massimino, Johnston, Noble, & Pearce-Higgins, 2015; Sullivan, Newson, & Pearce-Higgins, 2015) and our results show for which taxa regional analysis of biodiversity trends is likely to be feasible. Some taxa are reasonably-well recorded with the majority of species predicted to produce outputs above the precision threshold in many regions; these include pollinator groups such as bees and hoverflies that are the focus of much recent attention (Powney et al., 2019) and other popularly-recorded taxa. Our rules of thumb consider a standalone analysis of each regional dataset. This provides a conservative estimate of the suitability of the data, but this could be improved by conducting a joint analysis and allowing occupancy models to share information on detection probabilities across regions. By making the assumption that detection is consistent, it efficiently allows occupancy to be estimated individually for regions.

There is huge potential for volunteer-engagement and citizen science for biodiversity monitoring globally for a wide range of taxa. The availability of occurrence data is likely to rise through the increasingly popularity of tools for recording biodiversity sightings such as iNaturalist (https://www.inaturalist.org/), so that people can make records when and from where they choose. Our rules of thumb were developed from the specific case of biological recording in Great Britain, but they do provide guidance on sample size for analysis for citizen science multi-species recording projects for any taxa anywhere in the world. We also highlight the importance of considering the representativeness of the data and identify areas for further research in order to better support the design and implementation of these projects.

## Acknowledgements

None of this work would be possible without the incredible work of paid and voluntary organisers of recording schemes and the thousands of hours contributed by volunteer biological recorders across the UK each year. The Biological Records Centre (BRC) is organized and funded by the Centre for Ecology & Hydrology and the Joint Nature Conservation Committee. This work was supported by the Terrestrial Surveillance Development and Analysis partnership of the Centre for Ecology & Hydrology, British Trust for Ornithology and the Joint Nature Conservation Committee and by the Natural Environment Research Council award number NE/R016429/1 as part of the UK-SCAPE programme delivering National Capability. We thank Gary Powney for support and comments on the manuscript.

## Appendix S1: Characteristics of the datasets used in this analysis

**Table S1.**
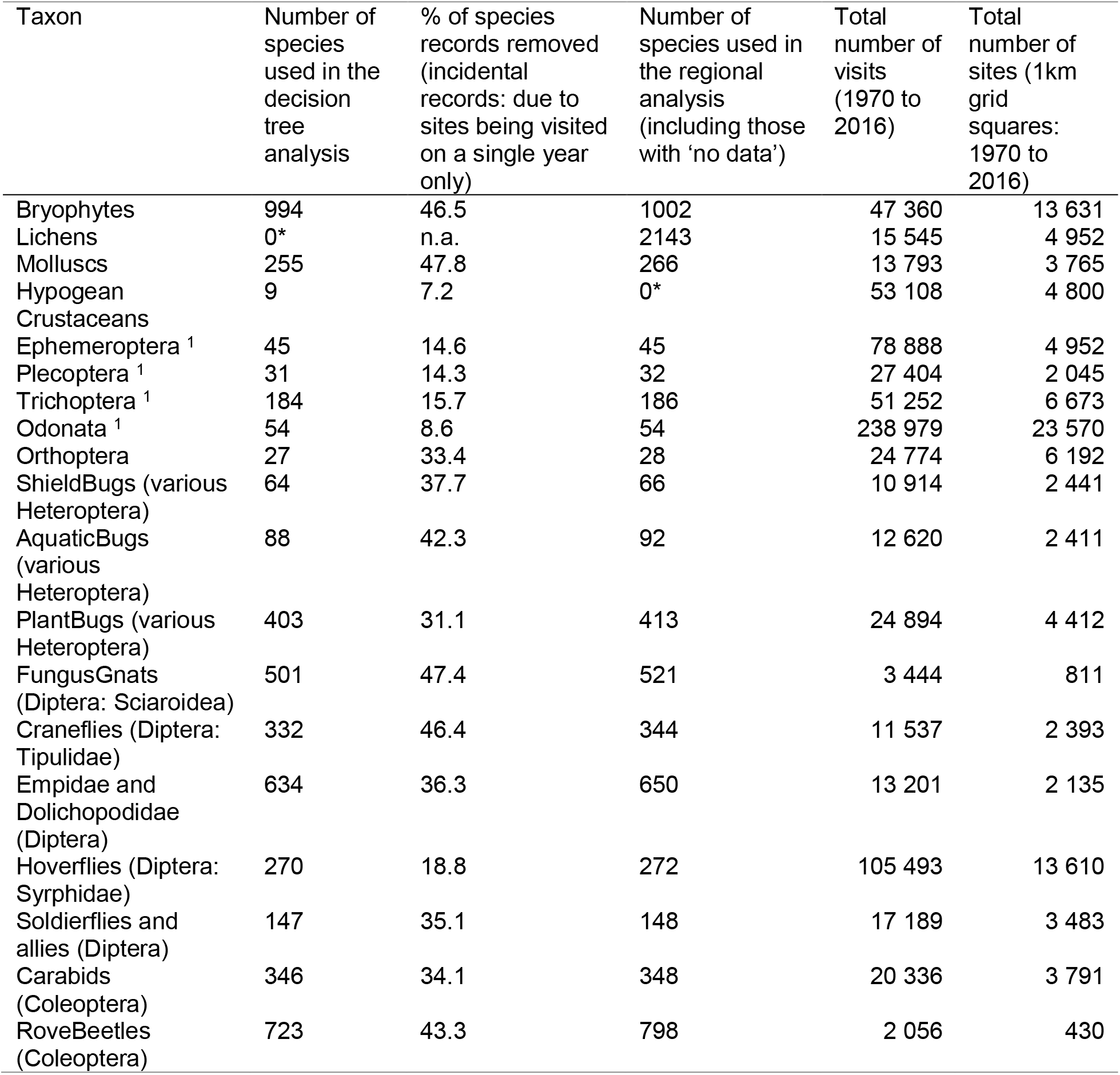
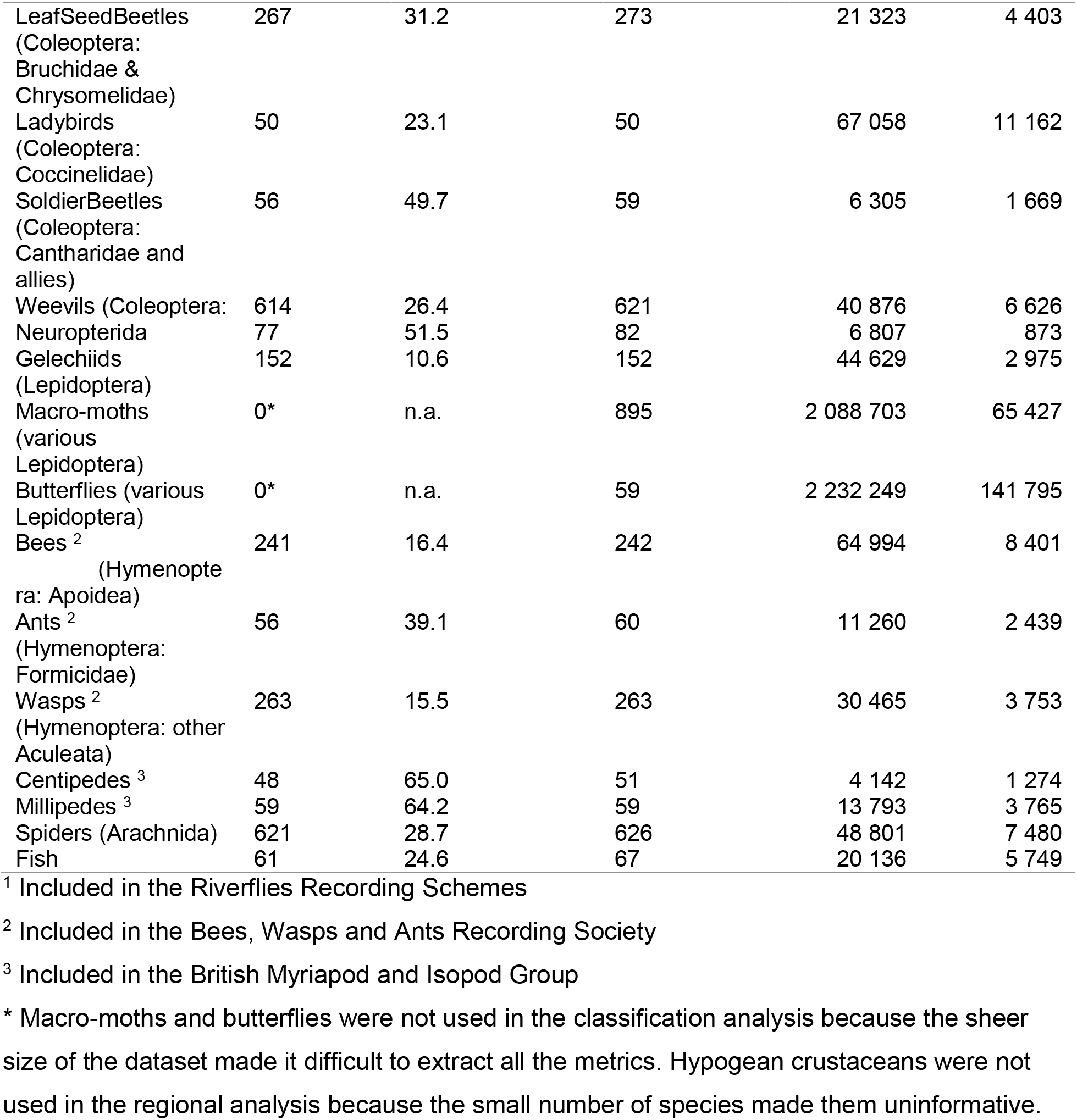
Number of species according to the individual projects. Taxonomic groups with the same superscript had data collated by the same recording scheme, but are taxonomically sufficiently different from each other that they were treated as separate projects in this analysis. Full details of the recording schemes are at https://www.brc.ac.uk/recording-schemes

**Fig. S1.1.**
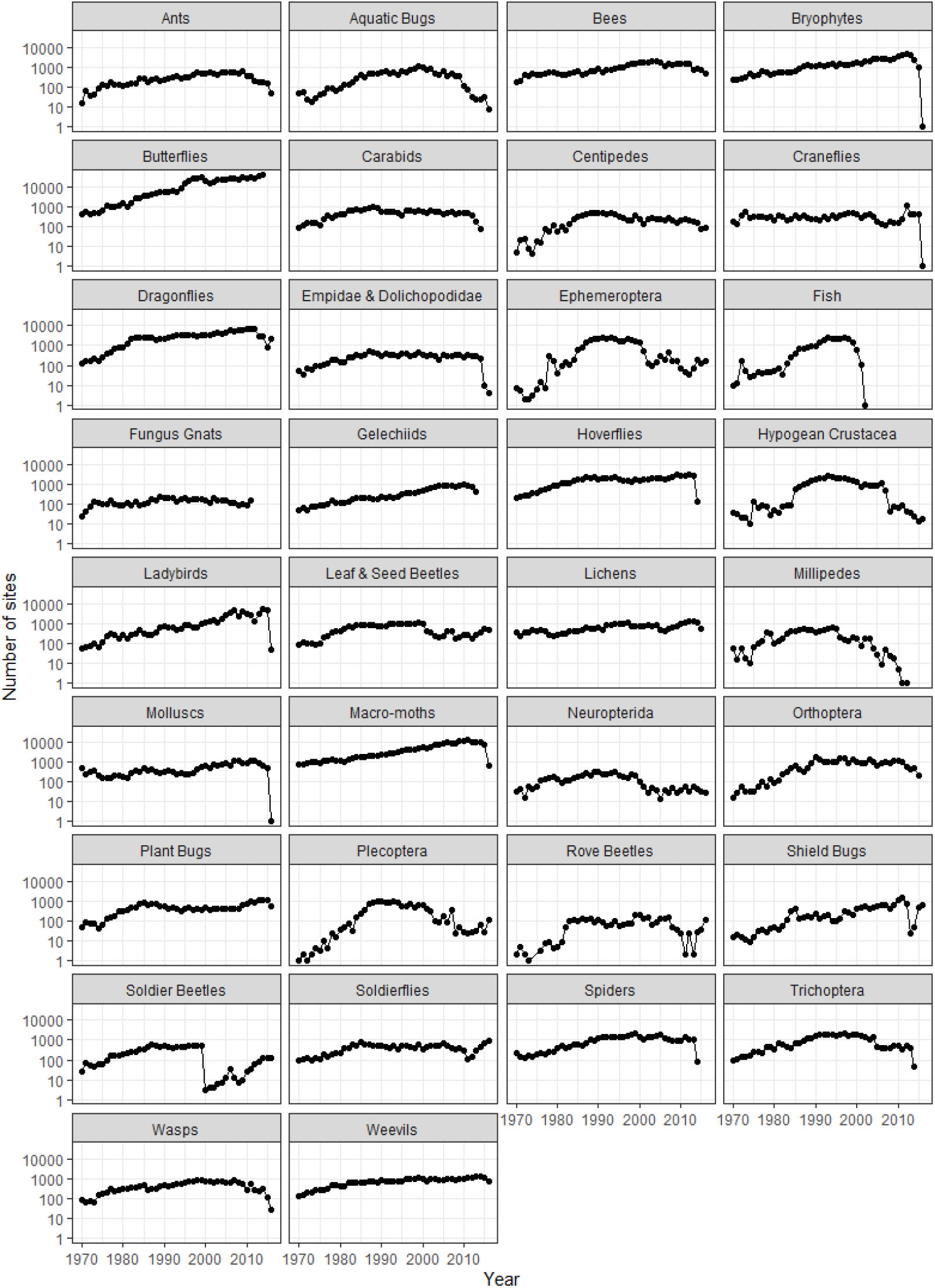
The number of sites (i.e. 1 km squares, on a log-scale) that have been visited varies substantially between taxa as shown by the summary across taxa of the median across time: median (quartiles) = 396 (180, 657) sites. For many groups there is a slight reduction in sites over the last one to five years due to delays in submitted data being verified and added to the databases.

**Fig. S1.2.**
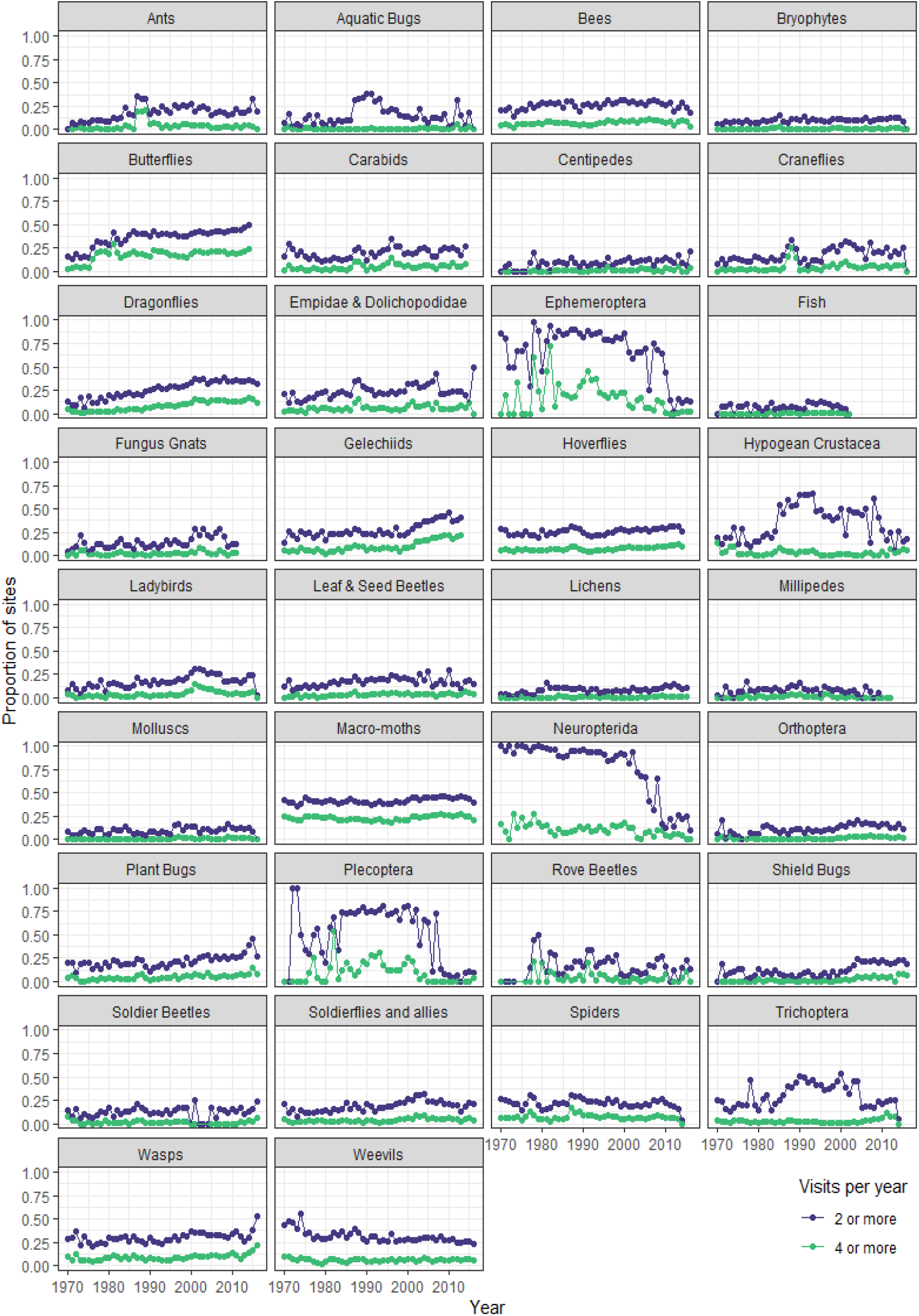
The proportion of sites visited in a year that were visited more than twice (black) or more than four times (grey) is fairly consistent for most taxa. The summary across taxa of the median across time: median (quartiles) proportion of sites visited two or more times = 0.20 (0.12, 0.27), and visited four or more times = 0.03 (0.01, 0.07). There are some taxa for which there is a high proportion of sites visited twice or more during the 1980s, 1990s and 2000s. Many groups are those found in freshwater, at least in their larval stages: aquatic bugs, Ephemeroptera, hypogean crustacea, Plecoptera, Trichoptera, and Megaloptera (within the Neuroptidera). This may be due to structured sampling for these taxa during surveys of freshwater quality.

## Appendix S2. Relationships between the data metrics

**Fig. S2.1.**
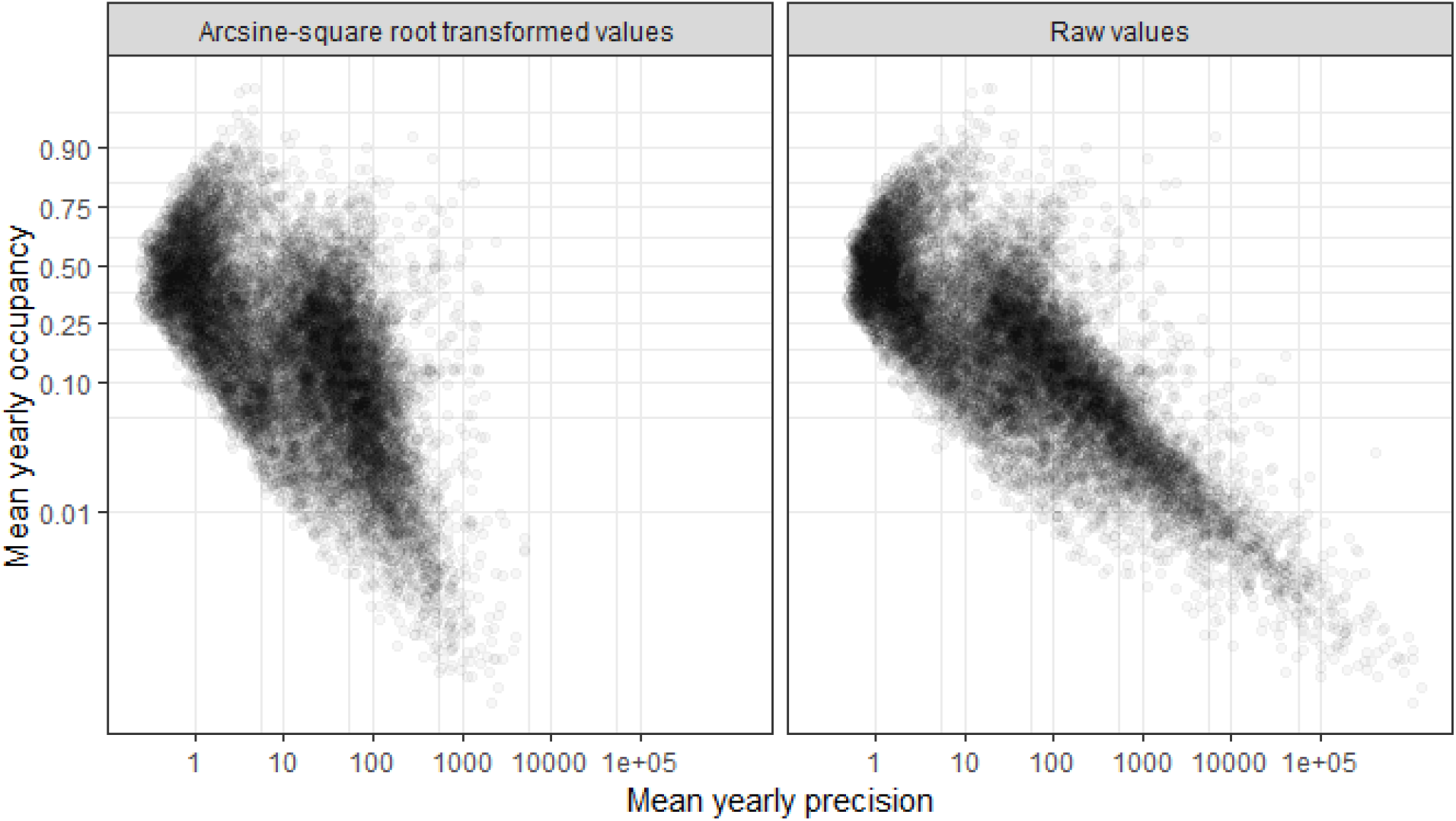
Comparison between the mean yearly precision for the raw posterior samples from the Bayesian occupancy model and the arcsine-transformed values. Note that for the raw values the precision is strongly related to mean yearly occupancy (outputs with low occupancy have very high precision, and trends with high precision are bound to be those with low mean yearly occupancy). This effect is lessened when the precision is calculated for the arcsine square root transformed values.

**Figure S2.**
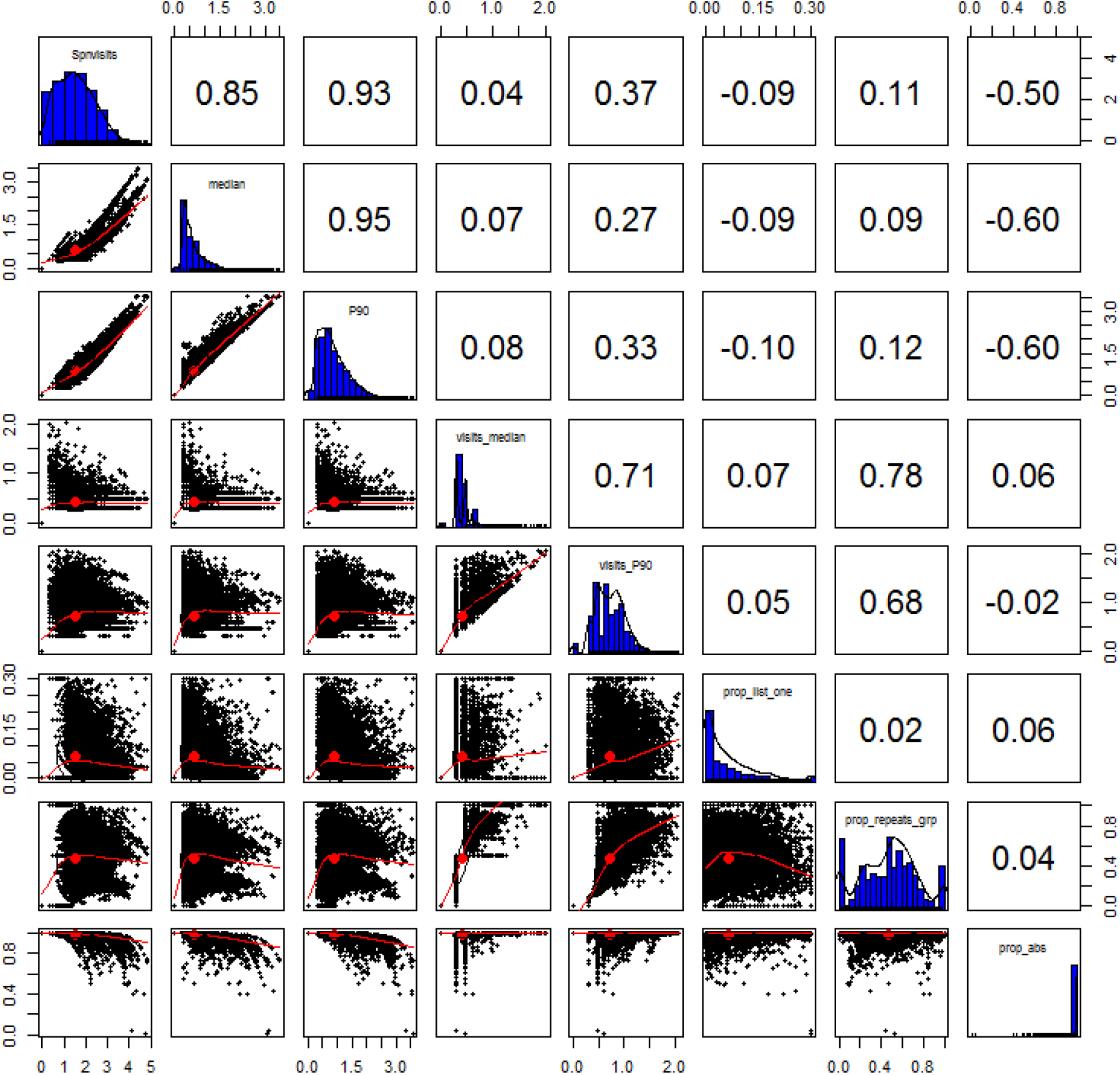
The associations between the total number of records (‘Spnvisits’) and the data metrics (Table 1 in the main text). Each dot represents one data output (a 47 or 10-year trend for a species, from the 31 taxonomic groups under consideration). Red lines show the smoothed relationship for ease of interpretation. Blue bars show the frequency distribution of the metric. Numbers above the diagonal show the correlation coefficient. Data metrics are: median = Median yearly number of records; P90 = 90th percentile yearly number of records; visits_median = Median yearly number of visits; visits_P90 = 90th percentile yearly number of visits; prop_list_one = Proportion of successful visits with list length one; prop_repeats_grp = Proportion of repeat visits for the group; prop_abs = Proportion of visits with non-detection of focal species). We used the transformation log10(X+1) for the metrics: median, P90, visits_median, visits_P90 and prop_list_one.

## Appendix S3. Further outputs from the decision trees

### Mixture model

We had no prior expectation that the frequency distribution of precision values would be bimodal, but it was notable that the precision threshold determined from expert elicitation (70.4) was close to the cross-over point of a Gaussian mixture model on the log10-transformed precision values (38.6). Only 7% of all the outputs had precision between these two values. The mixture model was run with ‘mixtools’ in R (Benaglia, Chauveau, Hunter, & Young, 2009).

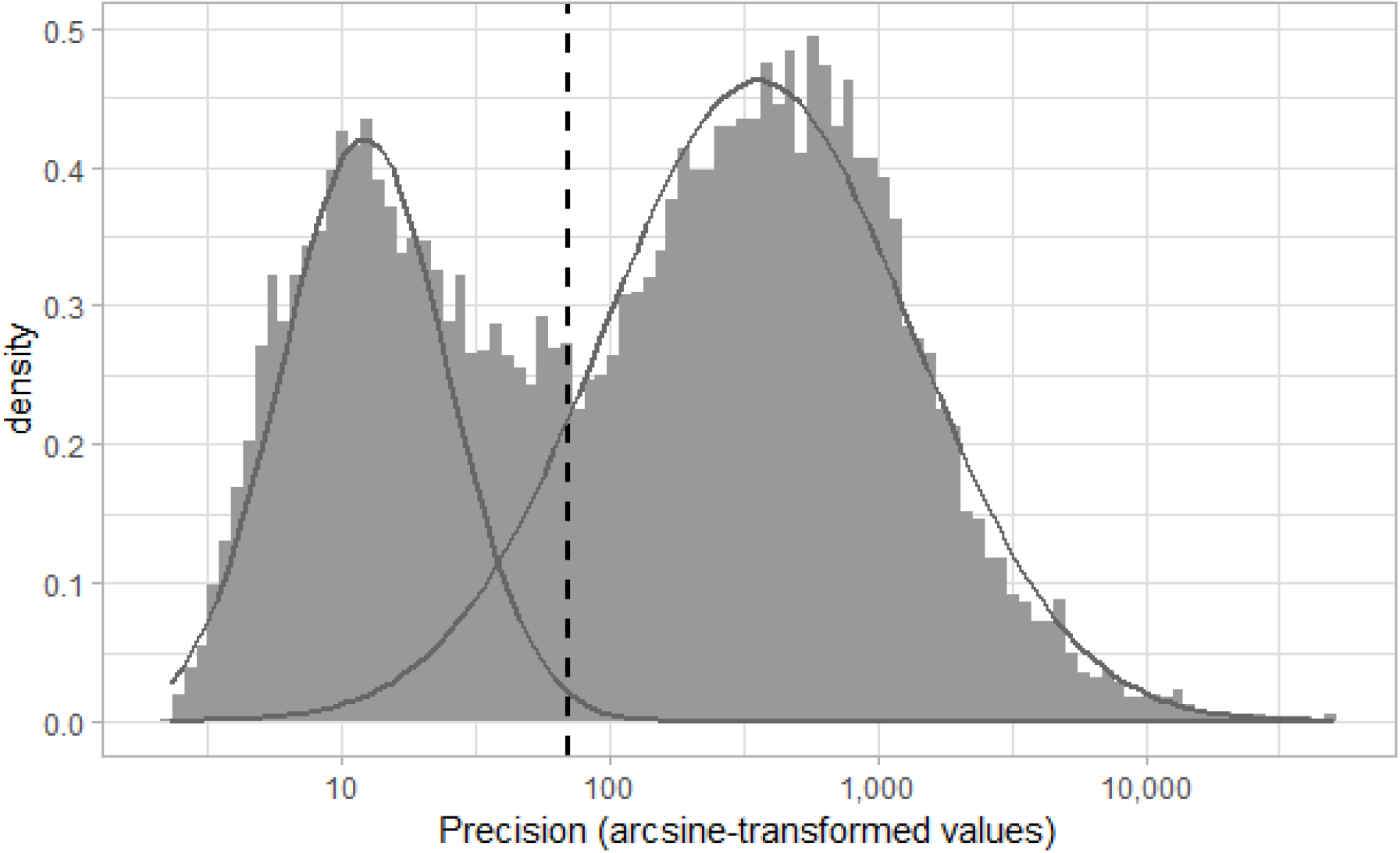

### Precision from untransformed versus arcsine-transformed data

**Fig. S3.1.**
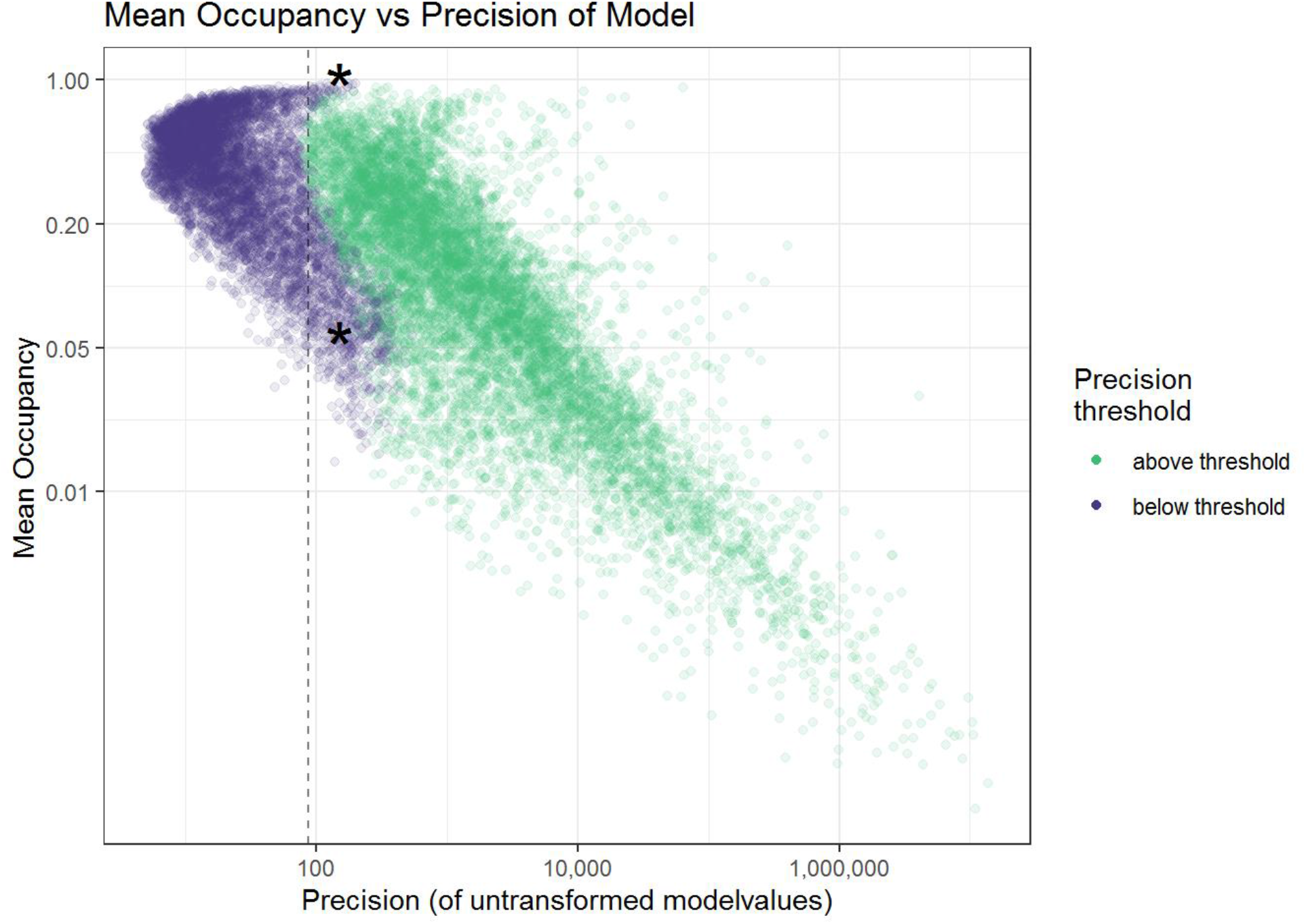
If we used untransformed posteriors from the Bayesian occupancy model outputs and repeated our analysis to define a precision threshold, then the precision threshold would be 87 (on the untransformed data). We found that there were a large number of outputs with high or low occupancy that were classified as above the threshold when using the untransformed data, but below when using the arcsine-transformed data (in the regions marked with * in the figure). The arcsine-transformation therefore reduces the boundary effect. This means that our precision threshold is defined so that if occupancy is close to 0 or 1, the precision (on untransformed data) needs to be higher for it to exceed the threshold.

**Table S3.1.** The classification statistics for the classification trees for acceptable precision and the two trees for high precision.

| Model | Specificity | Sensitivity | Accuracy |
| --- | --- | --- | --- |
| Equally-weighted | 79.4% | 85.7% | 83.0% |
| High specificity (best tree) | 98.7% | 49.2% | 70.2% |
| High specificity (matching equally-weighted) | 98.4% | 50.9% | 71.0% |

**Fig. S3.2.**
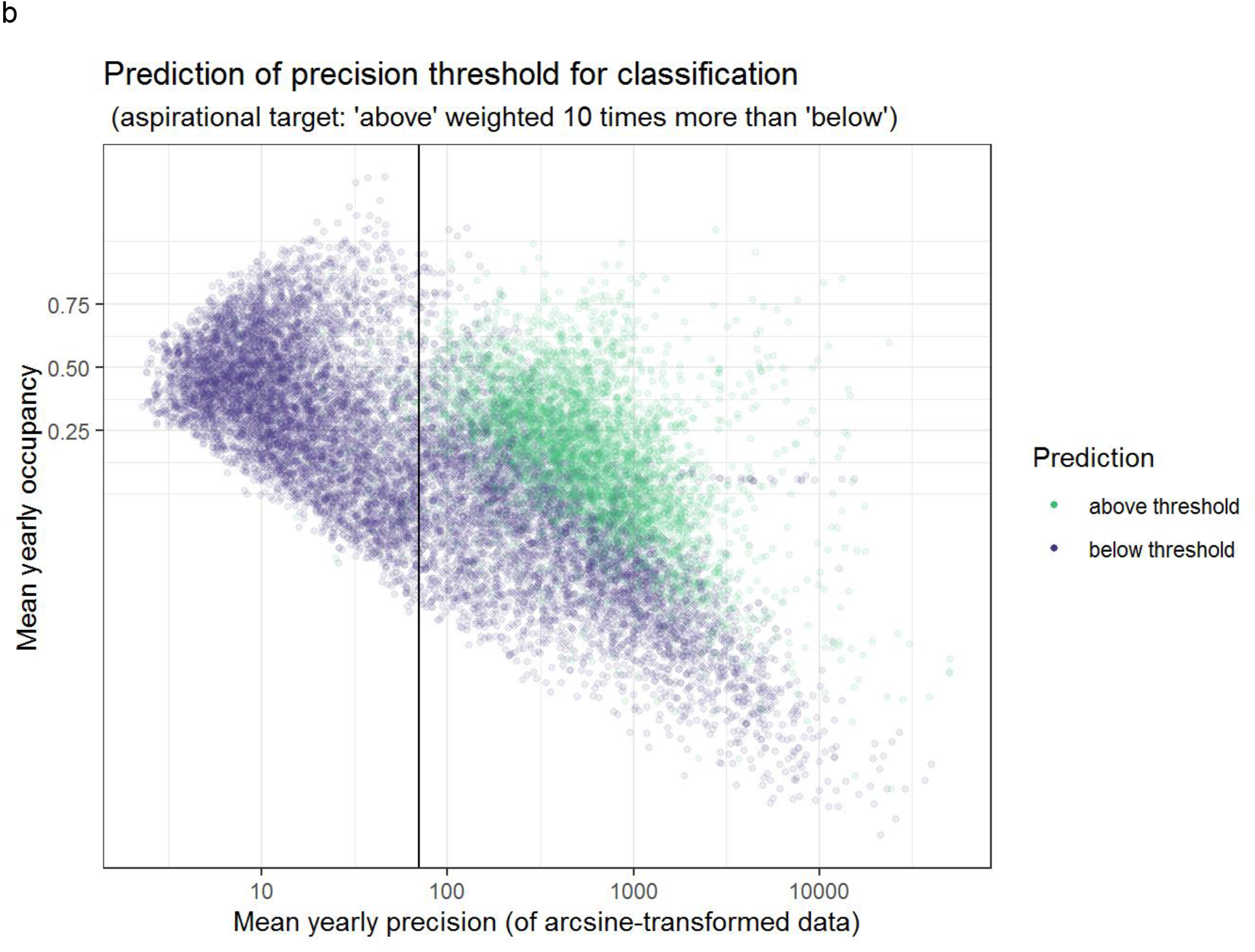
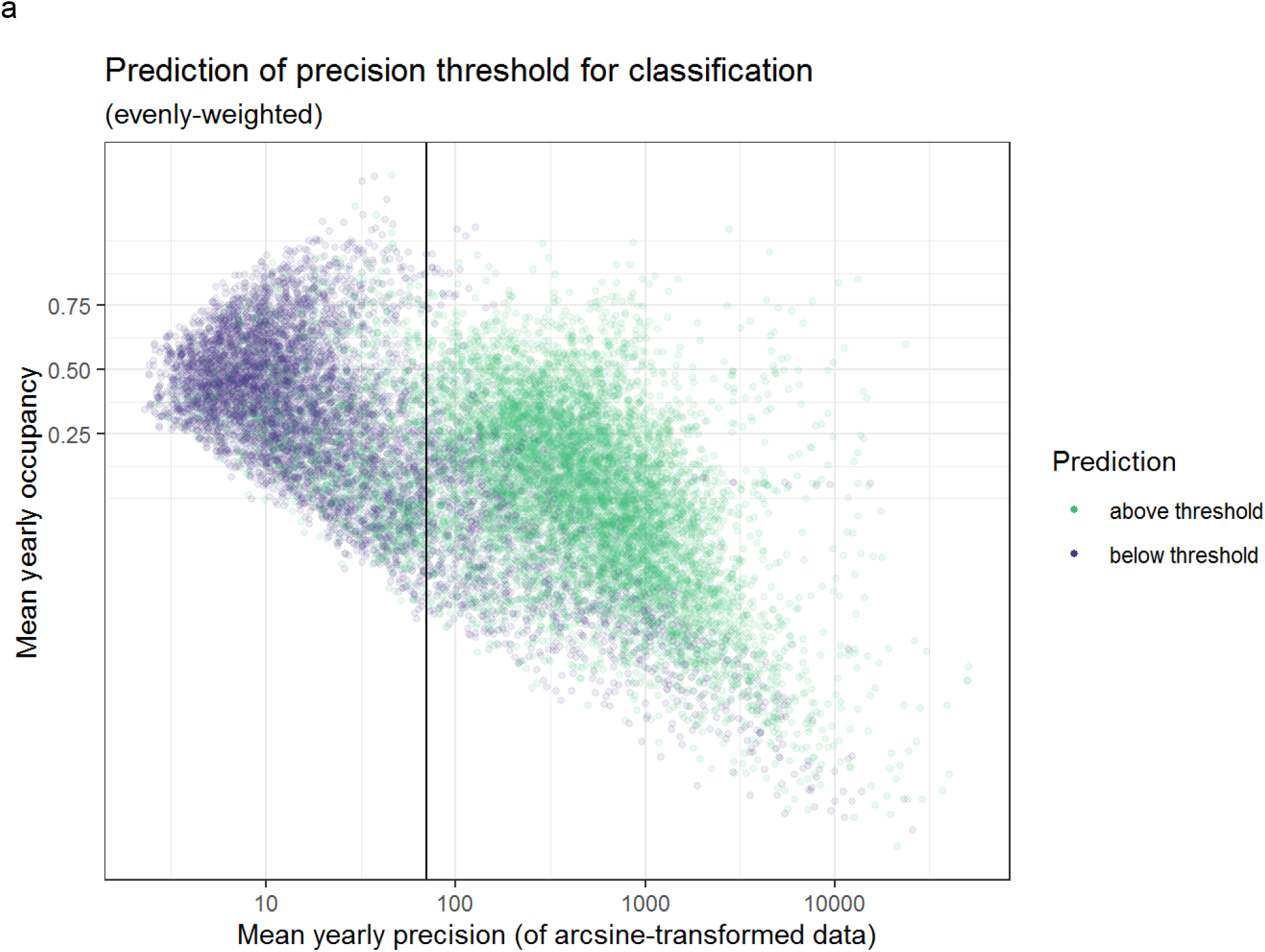
It can be seen that the aspirational decision tree mis-classified many more outputs, but any outputs that were classified as above the threshold (indicated by the black vertical line) were very likely to be truly above the threshold. This decision tree appeared to penalise the species with relatively lower estimated occupancy.

## Appendix S4

**Table S3.1.**
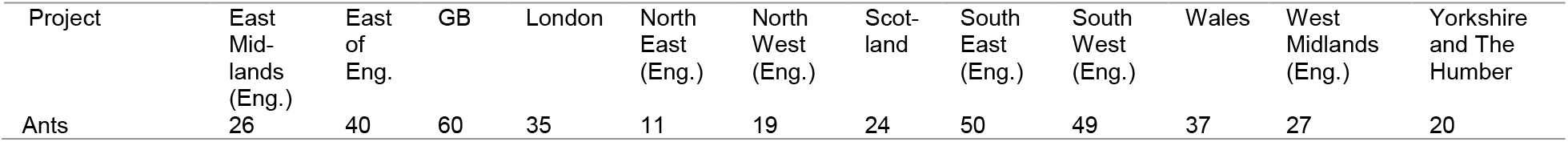
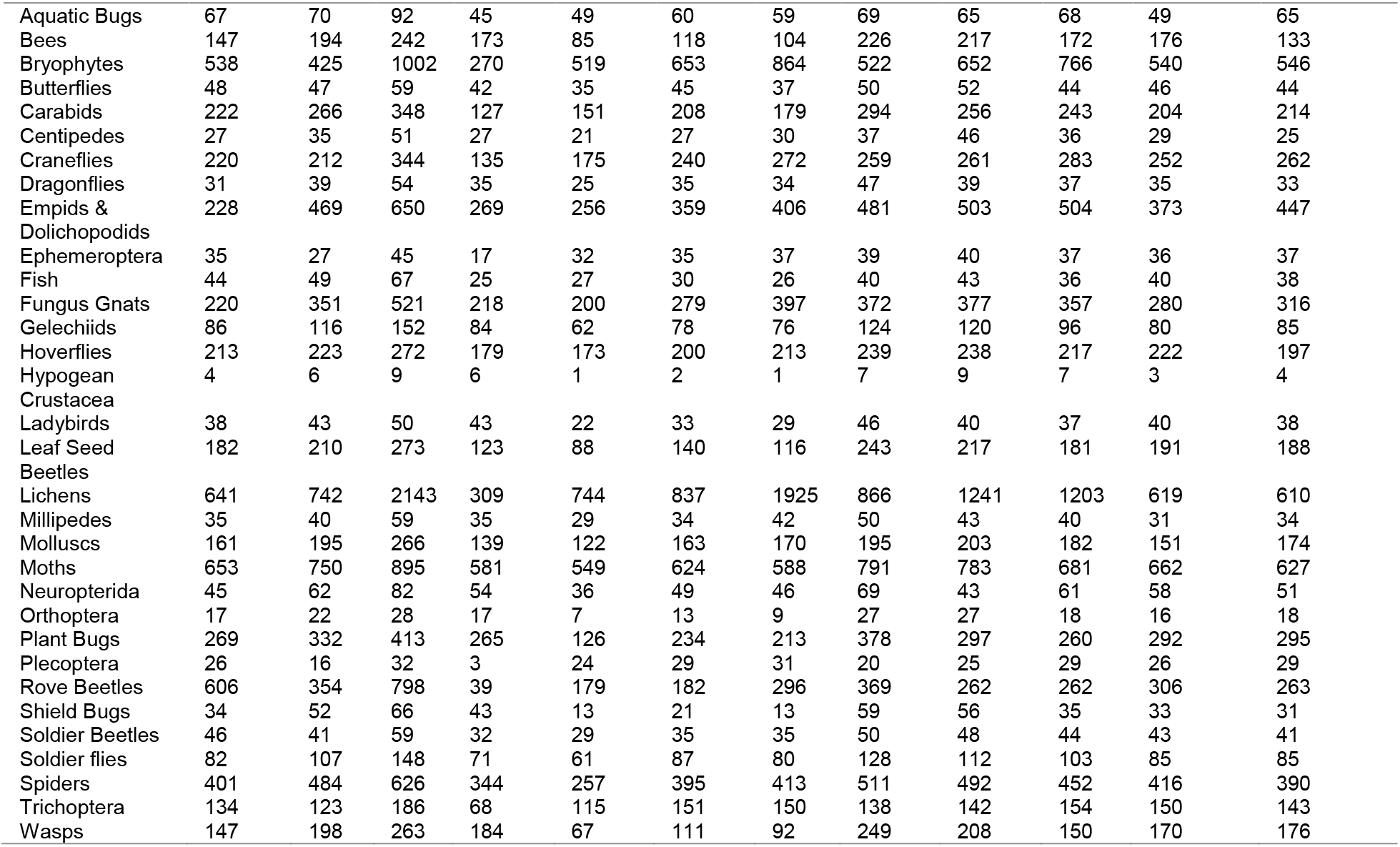
The total number of species for each taxonomic project in each region, as illustrated in Fig. 5 in the main text.

